# One size does not fit all: a novel approach for determining the Realised Viewshed Size for remote camera traps

**DOI:** 10.1101/2024.05.09.593241

**Authors:** B.M. Carswell, T. Avgar, G.M. Street, S.P. Boyle, E. Vander Wal

## Abstract

1. Camera traps (CTs) have become cemented as an important tool of wildlife research, yet, their utility is now extending beyond academics, as CTs can contribute to more inclusive place-based wildlife management. From advances in analytics and technology, CT-based density estimates of wildlife is an emerging field of research. Most CT-based density methods require an estimate of the size of the viewshed monitored by each CT, a parameter that may be highly variable and difficult to quantify.

2. Here, we developed and tested a standardized field and analytical method allowing us to predict the probability of photographic capture as it varies within CT viewshed. We investigated how capture probability changes due to environmental influences, i.e., vegetation structure, ambient temperature, speed of subject, time of day, in addition to internal factors from CTs themselves, i.e., sensitivity settings, number of photos taken, and CT brand. We then summarize these spatial capture probability kernels into a *Realised Viewshed Size* (RVS)—the capture-probability corrected size of a CTs viewshed

3. We found that RVS values are heavily influenced by location-specific environmental factors, i.e., vegetation structure, technological delays associated with CTs themselves, i.e., refractory period, and internal CT settings, i.e., sensitivity, number of photographs taken. We also found that the RVS values computed using our methodology are substantially smaller than reported values in the literature.

4. Imprecision surrounding CT viewshed areas can create propagating bias when implementing CT-based density estimators. Our method can change how practitioners consider photographs for use in CT density estimators thus increasing the reliability of CT-based density estimation, and contribute to more accessible wildlife management practices.

## Introduction

### Context

In the last five decades, camera traps (CTs) have become an established method of studying and monitoring wildlife populations (Fisher, 2023; Sollmann, 2018). CTs are an easy-to-operate, accessible, relatively affordable, and low-impact method of monitoring wildlife. CTs have become an integral part of wildlife management allowing researchers to answer location-based questions of occupancy (e.g., Tobler et al., 2015; Neilson et al., 2018), movement (e.g., Tape and Gustine, 2014), and behaviour (e.g., Caravaggi et al., 2017). Despite their common use, ambiguities around CT performance leads to uncertainty and decreasing the quality of inference made from some CT-based science. Among their limitations (e.g., Burton et al., 2015; Foster & Harmsen, 2012; Kolowski et al., 2021; Urbanek et al., 2019), empirically estimating the viewshed area a CT can monitor is an understudied, yet, critical facet when considering many CT-based analyses.

CT-based density and abundance estimation methods, henceforth referred to as viewshed density estimators (Moeller et al., 2023), are a novel group of statistical models incorporating different processes to estimate wildlife density (e.g., Rowcliffe et al., 2008; Moeller et al., 2018; Nakashima et al., 2020; Becker et al., 2022). Using CTs to estimate wildlife density promises to lower financial and logistic barriers currently in place for other traditional density estimation methods, e.g., aerial surveys. Some viewshed density estimators do not require the identification of individual animals and thus may constrain less, e.g., no need to mark individuals or census a population, than their traditional counterparts, such as physical mark-recapture methods or aerial flight surveys. Each viewshed density estimator, however, comes with its own set of assumptions and require novel parameters that we lack precision in estimating. For example, many viewshed density estimators require a precise estimate of the sampling area that CT’s monitor (Becker et al., 2022; Moeller et al., 2023; Nakashima et al., 2020; Rowcliffe et al., 2008; Rowcliffe et al. 2011).

Previous research has developed methods to quantify CT viewshed area (Table 1). Commonly used methods to estimate viewshed area include assuming a viewshed area or shape (Campos-Candela et al., 2018; Lyet et al., 2024; Figure 1), determining spatial bounds of CT viewshed (Cusack et al., 2015; Moeller et al., 2018; Figure 1A), and the Effective Detection Distance (EDD; Hofmeester et al., 2017; Howe et al., 2017; Rowcliffe et al., 2011; Figure 1B) that is based on distance sampling theory (Buckland, 2004). Each existing method to estimate viewshed area has its own unique application and set of limiting assumptions (Table 1). EDD for example, can account for obstructions in the viewshed and variation in study species size, however, the assumptions of distance sampling, i.e., perfect detection at 0 m (e.g., Figure 1B), are unrealistic as CTs cannot account for missed captures due to Passive Infrared (PIR) motion sensors. Additionally, methods to estimate EDD either require additional field apparatuses, i.e., marked distance poles, or require substantial *post-hoc* data processing and analysis, which are impractical in some geography and research programs (Haucke et al., 2022). Other methods, such determining viewshed bounds (Cusack et al., 2015; Moeller et al., 2018; Figure 1) or assuming manufacture determined areas (Campos-Candela et al., 2018; Lyet et al., 2024; Figure 1A) may be adequate for some CT studies, i.e., timelapse photography, yet, such methods assume perfect capture probability within determined bounds—unrealistic for PIR-based CT studies.

**Table 1.**
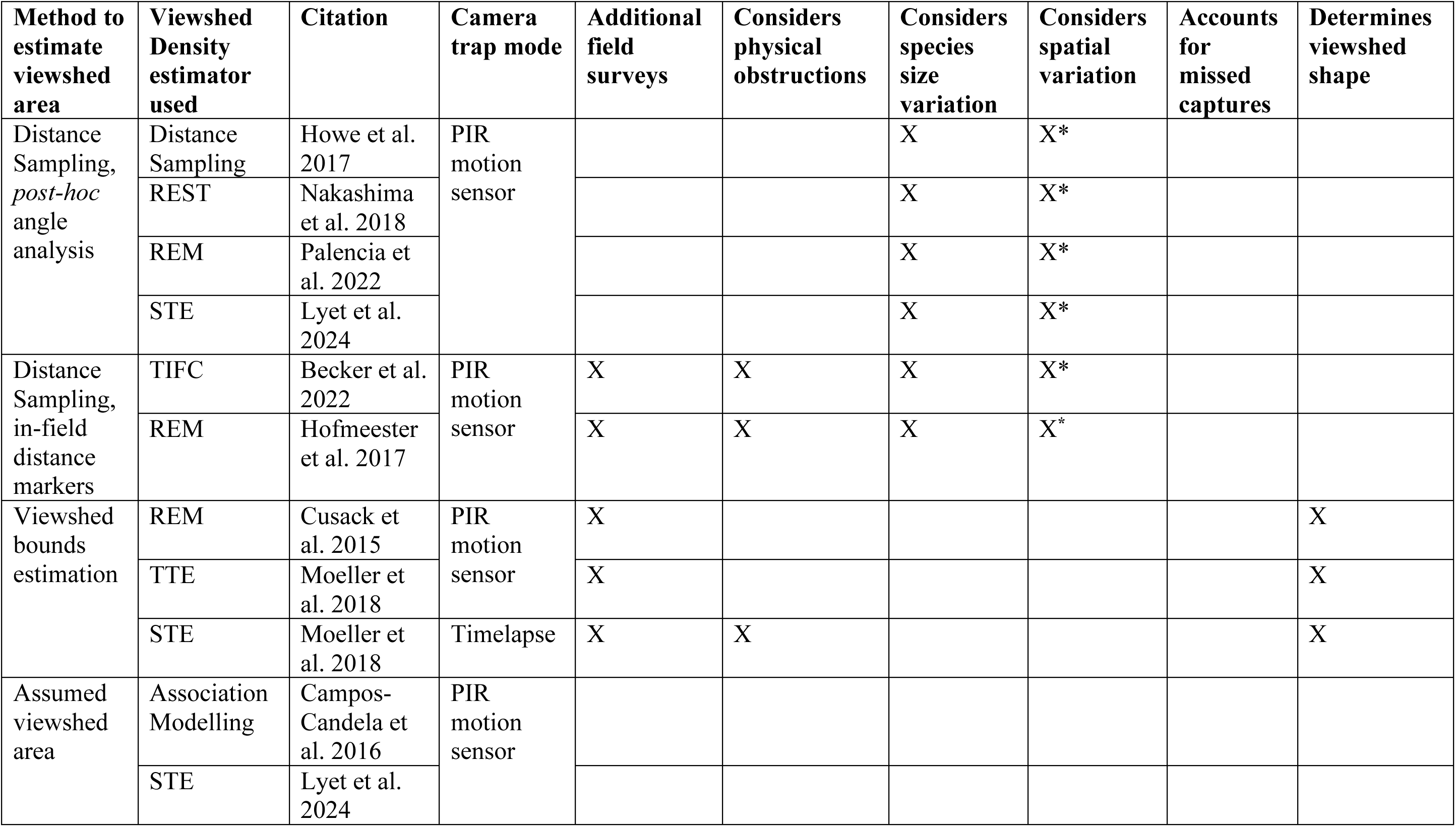
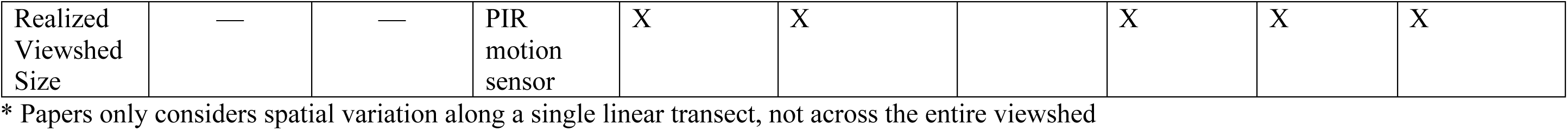
Commonly used methods to estimate viewshed area in camera trap literature, including which viewshed density model (i.e., Random Encounter Model [REM], Random Encounter Staying Time [REST], Time in Front of Camera [TIFC], Distance Sampling, Space-to-Event [STE], Time-to-Event [TTE]), and camera trap mode was used (i.e., Passive Infrared [PIR] motion sensor or timelapse photography), and various assumptions made about the viewshed area.

| Method to estimate viewshed area | Viewshed Density estimator used | Citation | Camera trap mode | Additional field surveys | Considers physical obstructions | Considers species size variation | Considers spatial variation | Accounts for missed captures | Determines viewshed shape |
| --- | --- | --- | --- | --- | --- | --- | --- | --- | --- |
| Distance Sampling, *post-hoc* angle analysis | Distance Sampling | Howe et al. 2017 | PIR motion sensor |  |  | X | X* |  |  |
|  | REST | Nakashima et al. 2018 |  |  |  | X | X* |  |  |
|  | REM | Palencia et al. 2022 |  |  |  | X | X* |  |  |
|  | STE | Lyet et al. 2024 |  |  |  | X | X* |  |  |
| Distance Sampling, in-field distance markers | TIFC | Becker et al. 2022 | PIR motion sensor | X | X | X | X* |  |  |
|  | REM | Hofmeester et al. 2017 |  | X | X | X | X* |  |  |
| Viewshed bounds estimation | REM | Cusack et al. 2015 | PIR motion sensor | X |  |  |  |  | X |
|  | TTE | Moeller et al. 2018 |  | X |  |  |  |  | X |
|  | STE | Moeller et al. 2018 | Timelapse | X | X |  |  |  | X |
| Assumed viewshed area | Association Modelling | Campos-Candela et al. 2016 | PIR motion sensor |  |  |  |  |  |  |
|  | STE | Lyet et al. 2024 |  |  |  |  |  |  |  |
| Realized<br>Viewshed<br>Size | — | — | PIR<br>motion<br>sensor | X | X |  | X | X | X |

\* Papers only considers spatial variation along a single linear transect, not across the entire viewshed

**Figure 1:**
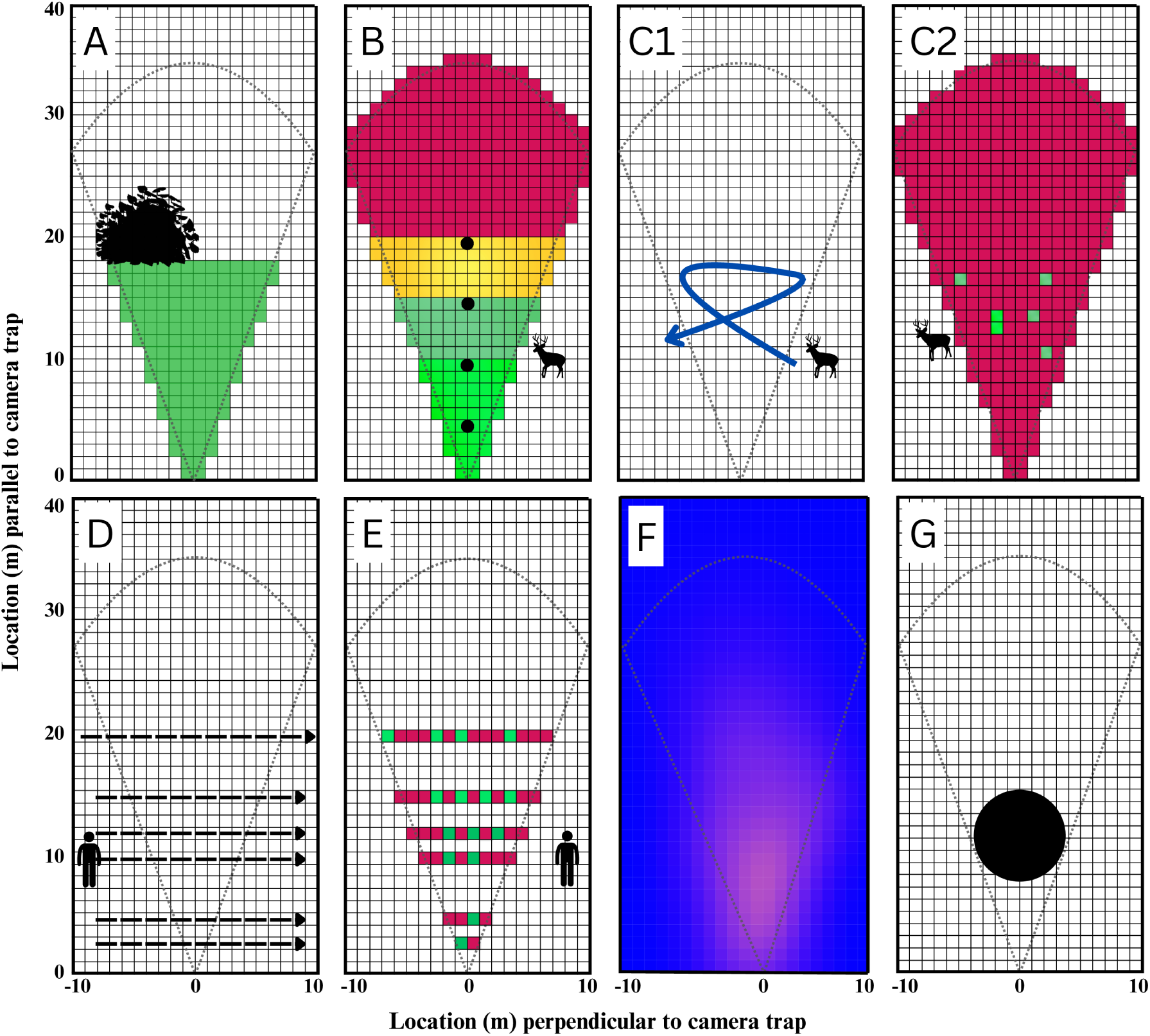
Various depictions of capture probability within camera trap (CT) viewshed; an assumed viewshed area based on manufacturer settings (gray cone; e.g., Campos-Candela et al., 2018; Lyet et al., 2024). **A.** Viewshed bound estimation based on physical obstructions—which assumes perfect capture probability within bounds (e.g., Cusack et al., 2015). **B.** Effective Detection Distance, based on in-field measuring poles, which considers the decay of capture probability with linear distance from the CT’s focal axis (e.g., Becker et al., 2022; Hofmeester et al., 2017; Rowcliffe et al., 2011). **(C1)** When an animal crosses the CT viewshed the number of photographs and thus capture probability spatially varies. The CT typically does not capture photos across all viewshed space for the time an animal is in the viewshed (green squares in **C2**). Our proposed method, which estimates the Realised Viewshed Size (RVS), uses jog-tests at fixed locations across the viewshed (**D**) to determine precisely where a subject was when photographs were taken or not taken (**E**). Using kernel-based analytics, we estimate capture probability accounting for missed captures and as it varies spatially across camera viewshed (**E**). We integrate spatially variable capture probabilities to represent the RVS—CT viewshed size corrected for perfect capture probability. The RVS is a theoretical value relevant for some viewshed density estimators (e.g., Random Encounter Staying Time, Time in Front of Camera models). Animal detections will occur outside of the theoretical RVS, but their probabilities are integrated into a minimum size with 100% capture probability.

### Focus

Imprecise estimates of the viewshed area are problematical because CTs, even when large number being used together, monitor a relatively small area—often a minute percentage of a study region. As a result, small errors in the determining the sampling area can propagate to large biases in abundance estimates when extrapolating fine-scale density estimates (Moeller et al., 2023). In addition, the theoretical area a CT monitors, as described by the CT’s manufacturer, does not align with the realised area. For example, Reconyx Hyperfire CT, a popular brand in wildlife research, market a viewshed area extending up to 30 m from the CT and 40° angle perpendicular to the CT’s lens, an area approximating 315 m^2^ (Reconyx, 2022; Figure 1). In reality, environmental factors such as habitat structure, topography, and vegetation (Moeller et al., 2023; Moll et al., 2020; Sultaire et al., 2023), will dictate a CT’s viewshed area for each unique location where CTs are placed (Apps and McNutt, 2018; Urbanek et al., 2019).

Researchers also need to account for the probability a photo will be taken given an animal is located within the CT’s viewshed, henceforth referred to as capture probability (Findlay et al., 2020; Moeller et al., 2023). Capture probability is more complicated and depends on numerous, interacting conditions caused by the environment in which CTs are placed, and CT settings. Capture probability is influenced by the CT’s Passive Infrared (PIR) motion detectors. PIR motion detectors require recognition of movement within pre-programmed zones (Urbanek et al., 2019), and a heat signature that contrasts ambient temperature for a photograph capture (Reconyx, 2022; Welbourne et al., 2016). PIR performance influences capture probability. Yet, few viewshed size estimation methods can adequately account for capture probability (Table 1).

Internal CT functioning and settings also dictate the viewshed area and how effectively CTs take photos (Apps & McNutt, 2018; Becker et al., 2022; Lepard et al., 2019; Urbanek et al., 2019). Even on rapid-fire modes, many CT user manuals report a 1–2 s refractory period from the time the PIR motion detector is triggered until that photograph can be written to the memory storage device (Del Bosco, 2021; Paula et al., 2014; Reconyx, 2022). Because a CT must reset and register another trigger for subsequent photos to be taken, this phenomenon would be sequential across the entire period an animal is in a CT’s viewshed. Thus, the trigger-to-photo delay can result in missed captures, i.e., when an animal is moving across the viewshed but no photographs are taken (e.g., Figure 1C1, 1C2). To decrease false triggers, e.g., from vegetation moving, researchers often lower CT sensitivity settings, influencing CT viewshed size. To aid in identification of species from photos, it is common practice to set CTs to capture multiple rapid-fire photos. Researchers often consider a photo series (i.e., all 3, 5, 10 photos) as a single event. As it takes CTs more time to capture more photographs, however, increasing the number of photos per trigger will reduce viewshed size. Effects of CT settings have not been empirically incorporated into CT literature, yet, have compounding influences on how effective CTs are at photographing wildlife.

Viewshed density estimators are often criticized for estimates that inconsistently align with traditional density estimation methods (e.g., Palencia et al., 2021; Becker et al., 2022; Fisher et al., 2023; Koetke et al., 2024). Although there may be many explanations for inconsistent results from viewshed density estimators, a lack of accuracy in, or entirely absent incorporation of viewshed area is undoubtedly one reason behind these discrepancies (Moeller et al., 2023). Here, we tested a standardized field protocol to 1) account for spatial variation and missed captures within the viewshed, 2) enumerate a kernel-based capture probability and how it changes with space in front of CTs, and 3) summarize capture probability kernels into a *Realised Viewshed Size* (RVS)—the capture probability corrected size of a CT’s viewshed.

## Methods

### Overall approach

The workflow we have developed, and detail below, is based on the idea that different locations in front of a CT will have differential capture probabilities due to factors such as position relative to the PIR sensor, vegetation, and microterrain between the location and the sensor. In general, we expect spatially variable capture probabilities to be highest near the PIR’s focal point (along the optical, at a distance of 10 meters), and decay to 0 at prereferral locations. We recognize that the resulting spatial capture probability surface may be complex, with local valleys and peaks, and vary substantially from one deployment to another. Our aim was thus to develop a workflow that enables us to quantify this spatial capture probability surface, such that, we can then integrate over the capture probability surface to obtain the Realised Viewshed Size (RVS)—the denominator of the Random Encounter Staying Time (REST)/ Time in Front of Camera (TIFC) density estimator.

To estimate RVS, we present the CT with PIR triggering events (a person jogging) and analyse the resulting successful vs failed capture events, i.e., if a photograph was captured, as a function of the spatial coordinates of an event, resulting in an empirically derived spatial capture probability surface.

### Field methods

To assess capture probability, we tested a standardized field method in two separate case studies. We applied our methods to each CT to calculate capture probability, unique to each CT in our study that in Case Study A) accounts for differences due to habitat and other location specific influences and in Case Study B) incorporates variation resulting from internal CT settings. Adapting methods from other researchers (Apps & McNutt, 2018; Del Bosco, 2021; Palencia et al., 2021), we established six 20 m long transects, perpendicular to and centred at the CT’s optical axis at distances of 3, 5, 10, 12, 15, and 20 m from the lens (Figure 1D). These transects extended past CTs’ viewshed and all cameras were deployed at a height of ∼1.5 m. We performed a ‘jog-test’ where one researcher jogged at a speed of ∼2 m/s along each transect, six times, for a total of 36 jogs per CT. With a timer, another researcher initiated each jog-test by triggering the CT using a hand motion to take a photo and recorded the time in seconds it took to jog across the full 20 m. CTs captured PIR motion-triggered photos during jogs, just as they would when triggered by a moving animals, however, in our tests, we know exactly the location of the subject across space and time, and can hence confidently identify failed capture events to be contrasted with successful capture events.

### Analytical methods

To calculate capture probability, we determined the time elapsed (s) between the transect start times and the time when each photograph was taken. Next, we calculated the speed of each jog as the time taken to cross each transect divided by transect distance (m/s). We calculated spatial locations where successful capture events or ‘used locations’ occurred, by multiplying the time elapsed at each photograph by jog velocity to calculate the distance travelled along the transect, rounding to nearest meter and second. For trials where more than one photo were taken per trigger, we filtered photo series down to the first instance when a photograph of the jogger was taken. We filtered photo series down to one event to represent how many practitioners consider CT data. By using jog velocities, and transect start and end time, we calculated all possible spatial locations where photographs could have occurred or ‘available locations’. We then excluded ‘used’ locations from the set of ‘available locations, leaving us with a set of unused locations, i.e., failed capture events. Combined, the ‘used’ and ‘unused’ locations were projected onto a 1 m grid-system along each transect where each cell had a Bernoulli outcome, whether a photographic capture occurred or not (Figure 1E).

### Statistical models

We fit Bernoulli capture data using Generalized Additive Mixed Models (GAMMs) using the mgcv package in Program R (Wood, 2017) for each case study, separately. We used GAMMs to estimate a spatial capture probability surface by fitting a Gaussian Process spline (i.e., Kriging) to the spatial coordinates of our used/unused data (Wood, 2017).

We used the resulting GAMM models, along with covariate data for each case study, and the *predict_gamm()* function (Wood, 2023) to predict the probability of photographic capture occurring across a 1×1 m spatial grid—a discrete approximation of the undelaying capture probability surface (Figure 1F). We further applied these models to also predict capture probability at any CTs where data were collected, e.g., vegetation cover, ambient temperature, but that were not included in the trial tests.

Our primary goal was to determine the size of the theoretical area that a CT monitors with 100% capture probability: the RVS. To estimate the RVS, we summed the GAMM-based probability of photographic capture across each 1 m^2^ grid cell with a rectangular area 20 m wide and 40 m long (800 grid cells; see Figure 1G), resulting in an RVS estimator in units of m^2^. Our approach assumes that the capture probability outside of 20 x 40 m rectangle is negligibly small, which is likely the case for most CT designs. Our approach further assumes equivalency between predicted probabilities of photographic capture and the percent area monitored in each cell. This assumption of equivalency is a fundamental assumption underlying other, popular analyses for CT data such as occupancy modelling (MacKenzie et al., 2002). Capture probabilities represent the equivalent area within each cell that a photograph would be captured with certainty. For example, a capture probability of 0.33 in a 1 m^2^ cell is equivalent to having 0.33 m^2^ of area where a capture will occur 100% of the time. In addition, we determined the uncertainty associated with each RVS calculation.

We wanted to ensure sufficient predictive accuracy of our analysis for our Case Study A field trial. Because a CT’s monitoring area is unknown, we employed *k*-folds cross validation on the probability of photographic capture for spatial locations in front of each CT (Geisser, 1975). Using unique CT ID as a blocking variable, we withheld entire CTs to assess the predictive accuracy of each 1×1-m cell across CT and thus the validity of our method at CTs where jog-tests were not conducted (Fielding & Bell, 1997). We generated Receiver Operating Characteristics (ROC) Area Under Curve (AUC) scores for each of these validations.

### Case Study A

In Case Study A, we tested 45 CTs actively deployed as a part of a long-term monitoring program located on Treaty 2 territory, the traditional lands of the Anishinabewaki, Očhéthi Šakówiŋ, Cree, Oji-Cree, and the Homeland of the Métis peoples (Riding Mountain National Park, Canada; RMNP; 50.658524, -99.970601). We conducted jog-tests on two different CT brands deployed at our research sites: Reconyx Ultrafire (*n* = 9; Reconyx, Holmen, USA) and Cuddeback H-1453 (*n* = 36; Cuddeback, Green Bay, USA). We did not vary any internal settings between trials in Case Study A—all CTs were set to a high sensitivity setting and a single photograph taken for each trigger. Instead, we measured external factors known to affect capture probability. Factors included: vegetation cover in front of CTs (Moeller et al., 2023; Moll et al., 2020), ambient temperature (°C) at time of survey (Urbanek et al., 2019), speed (m/s) of jog for each transect (McIntyre et al., 2020), and CT model (Apps and McNutt, 2018). In addition, to account for a refractory period after a photographic capture (Del Bosco, 2021; Paula et al., 2014; Reconyx, 2022), we recorded whether CTs captured a photo in the previous two seconds—we expected a negative effect of a recent capture on the probability of a current capture. To measure vegetation cover, we conducted two surveys: (1) a shrub cover survey counting multi-stemmed woody species 10 m in front of each CT by using the Daubenmire method on a 4 m radius plot (Daubenmire, 1959); and, (2) horizontal cover via a cover pole, placed 10 m in front of CTs to quantify vegetation height. For Case Study A, we created a single, saturated model including covariates measured in the field, i.e., parallel and perpendicular distances from CT, percent shrub, and vegetation cover in front of CT, ambient temperature (°C), velocity (m/s) of each jog, CT model (categorical), and whether a photograph had occurred in the previous 2 seconds (binary). Additionally, to allow different CTs to differ in their RVS above and beyond the effect of covariates, we implemented unique CT ID as a random factor smoother (Wood, 2017, 2023).

### Case Study B

For Case Study B, we conducted a series of jog-tests in an open, non-vegetated, flat field in a local park located on traditional unceded territories of the Beothuk and Mi’kma’ki peoples (St. John’s, Canada; 47.595254, -52.745665). The trial used a controlled habitat to isolate the effect of internal CT settings on capture probability. In total, we surveyed 27 Reconyx Hyperfire 2 (Reconyx, Holem, USA) CTs across 3 trials: two during daylight (∼14:00 h; during partially cloudy conditions) and one after sunset (∼23:00 h). These trials were conducted to measure the influences of the internal CT sensitivity settings (Heiniger & Gillespie, 2018), the number of photos taken per motion trigger and refractory period following a photograph capture (Paula et al., 2014). In addition, we wanted to measure the deterioration of CT performance during the nighttime. We held CT settings constant across trials using specific combinations of settings (see Table 2 for all combinations). Due to the lack of visual obstructions during the daytime trials in the open field, we observed a higher-than-expected capture probability on the 20 m transect.

**Table 2.** The combinations of camera sensitivity settings, number of photos taken for each passive infrared motion trigger, and time of day we implemented on to Reconyx Hyperfire 2 (*n* = 26) camera traps during Case Study B, in a controlled, open field setting, in St. John’s Canada.

| Trial type | Number of cameras | Sensitivity setting | Number of photos per trigger | Time of day |
| --- | --- | --- | --- | --- |
| Sensitivity | 3 | Low | 1 | Day |
|  | 3 | Medium | 1 | Day |
|  | 3 | Very high | 1 | Day |
| Number of photos | 3 | High | 1 | Day |
|  | 3 | High | 3 | Day |
|  | 3 | High | 5 | Day |
| Night | 9 | High | 1 | Night |

Thus, for the two daytime trials we jogged an additional two transects at 30 and 40 m parallel to CTs to fully enumerate the decay of capture probability with distance. For CTs assessed in Case Study B, in addition to the smoothed parallel and perpendicular locations (McIntyre et al., 2020), we included categorical covariates for each of the variables of interest: CT sensitivity setting, number of photos taken per motion trigger, time of day, and whether a capture had been taken in the previous two seconds (Table 3).

**Table 3.** Binary Generalized Additive Mixed Models (GAMMs) used in determining the Effective Capture Areas of (a) camera traps (*n* = 46) in Riding Mountain National Park, Canada and (b) camera traps (*n* = 27) in a non-vegetated local park near St. John’s, Canada. Showing different predictor variables used in each model, with their respective explanations, coefficient values, and probability values. Both models contain the parallel and perpendicular locations in front of each camera trap with splines implemented, in addition to unique camera identification implemented as a random intercept.

| Riding Mountain National Park field data (Case Study A) model: |  |  |  | Controlled setting data (Case Study B) model: |  |  |  |
| --- | --- | --- | --- | --- | --- | --- | --- |
| Covariate | Explanation | Coefficient<br>($\chi^2$ for<br>smoothed<br>terms) | p-value | Covariate | Explanation | Coefficient<br>($\chi^2$ for<br>smoothed<br>terms) | p-value |
| **s(parallel distance, perpendicular distance)** | Gaussian process spline of the parallel (locations in front) and perpendicular (locations to the side) of camera traps | **829.978** | **<0.001** | **s(parallel distance, perpendicular distance)** | Gaussian process spline on parallel (in front) and perpendicular locations (to the side) of camera traps | **1009.3** | **<0.001** |
| te(shrub cover, horizontal cover) | Tensor product spline of percent shrub cover (determined by Daubenmire plots) and horizontal cover (determined from cover pole) | 1.435 | 0.707 | Low camera sensitivity | Categorical variable for the sensitivity setting chosen on each camera trap for a specific trial | Set as intercept |  |
|  |  |  |  | **Medium camera sensitivity** |  | **1.076** | **<0.001** |
|  |  |  |  | **High camera sensitivity** |  | **1.837** | **<0.001** |
|  |  |  |  | **Very high camera sensitivity** |  | **3.479** | **<0.001** |
| **Photo taken in previous 2 seconds** | Binary variable of whether the camera took a photo in the previous two seconds | **-1.064** | **<0.001** | 1 photo per trigger | Categorical variable for the number of photos taken per trigger setting chosen on each camera trap for a specific trial | Set as intercept |  |
|  |  |  |  | 3 photos per trigger |  | **-1.266** | **<0.001** |
|  |  |  |  | 5 photos per trigger |  | **-1.186** | **<0.001** |
| Jog velocity |  | 0.211 | 0.126 | Nighttime trial |  | Set as intercept |  |
|  | Velocity of each<br>transect jog during<br>the field surveys |  |  | Day-time trial | Whether or not<br>trials occurred<br>during daylight<br>or night hours | **1.072** | **<0.001** |
| Ambient<br>temperature | Ambient<br>temperature (°C)<br>measured from<br>camera traps at time<br>of survey | 0.082 | 0.428 | **Photo taken in**<br>**previous 2**<br>**seconds** | Binary variable<br>of whether the<br>camera took a<br>photo in the<br>previous two<br>seconds | **0.173** | **0.047** |
| Camera model | Categorical variable<br>for the brand of<br>camera trap<br>(Reconyx or<br>cuddeback) used in<br>the field trials | -0.370 | 0.527 |  |  |  |  |
| **s(unique**<br>**camera ID)** | Random factor<br>smooth<br>implemented on<br>each unique camera<br>trap | **358.793** | **<0.001** | **s(unique**<br>**camera ID)** | Random factor<br>smooth<br>implemented on<br>each unique<br>camera trap | **24.7** | **<0.001** |

To determine how strongly numeric covariates in Case Study A influence a CT’s RVS, we calculated a *post-hoc* effect size through prediction. We ran separate predictions for each numeric covariate of interest—percent shrub cover, percent horizontal cover, velocity of jog, and ambient temperature—and generated RVS values across the observed range of each covariate while holding all other variables at their mean values. For example, horizontal cover ranged from 42–99%, thus we ran an RVS prediction for each unit (percent) value in that range. Next, we calculated the mean of all unit-change differences in RVS predictions. The resulting values represent the mean influence of a 1-unit change by a covariate on a CT’s RVS. For Case Study B, we ran similar predictions, but for each level in the categories of interest, i.e., sensitivity setting and number of photos per trigger, while holding all other variables at constant values.

## Results

### Case Study A

We determined the predicted probability of a successful photographic capture occurring within a CT viewshed at a 1 x 1 m resolution (Figure 2). For all CTs where we collected local site data, we calculated unique capture probability distributions and scaled them to a RVS. For example, Figure 2 represents a Reconyx Ultrafire CT within our Riding Mountain National Park field site, with an average 62% shrub cover, 79% horizontal cover, and an ambient air temperature of 1.5°C during survey time. Assuming a subject moving at a velocity of 2 m/s in the CTs viewshed area, we scaled these probabilities to a RVS of 15 m^2^ and a standard error ranging from 8–27 m^2^ (Figure 2). Ambient temperature at time of jog-test surveys ranged from -2–10°C, velocity of each jog from 0.76–2.86 m/s, percent shrub cover from 0–176%, and percent horizontal cover from 42–98%.

**Figure 2.**
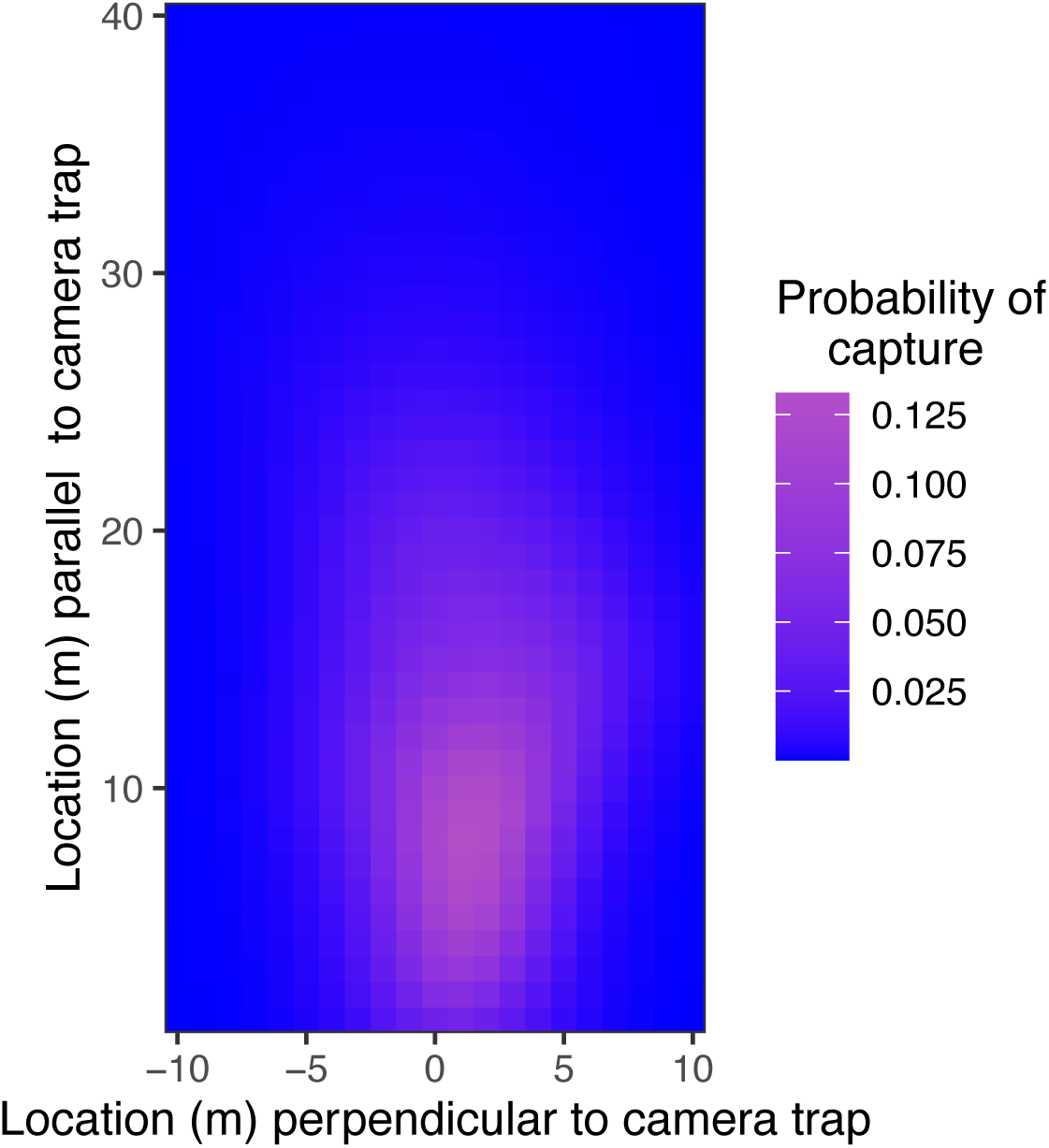
Predicted probabilities of photographic capture, determined by Generalized Additive Mixed Models, on a 1×1 m grid for an average Reconyx Ultrafire camera in Case Study A, with 62 % shrub cover, a subject moving at a 2 m/s velocity, and an ambient air temperature of 1.5°C. The predicted Realised Viewshed Size (± 1 SE) for this camera was determined to be 15 m^2^ (8– 27m^2^). The camera traps is located at position [0,0] on the grid.

Covariates in the model showed varying levels of significance (Table 3). The effect of spatial location in front of the CT, fitted as a Kriged surface using a Gaussian process spline, was highly significant (*F* = 42.655, *p* <0.001). In addition, the binary refractory period variable showed a significant negative relationship with photographic capture (*β* = -0.955, *p* <0.001).

Other covariates showed varying effects on the resulting RVS value (see Table 3). Percent unit changes in shrub (^x̅^ = 0.14 m^2^) and horizontal cover (^x̅^ = 0.62 m^2^) had a relatively small influence on RVS. Maximal effects between the largest and smallest RVS predictions, however, were larger. The difference between the largest and smallest RVS was 24.48 m^2^ for shrub cover and 35.60 m^2^ for horizontal cover (Figure 3). A unit change in temperature (°C) had a larger influence on RVS, with a mean of 3.11 m^2^, and a maximum of 37.32 m^2^ difference between the maximum and minimum RVS predictions (Figure 3). Finally, unit changes in velocity (m/s) of each jog had the largest influence on RVS prediction, with a mean of 6.97 m^2^ and a maximum of 13.95 m^2^ between the largest and smallest predictions (Figure 3). The cross validated predictive accuracy of the model, assessed at each 1×1 m cell between CTs, was good (AUC=0.76; Boyce et al., 2002).

**Figure 3.**
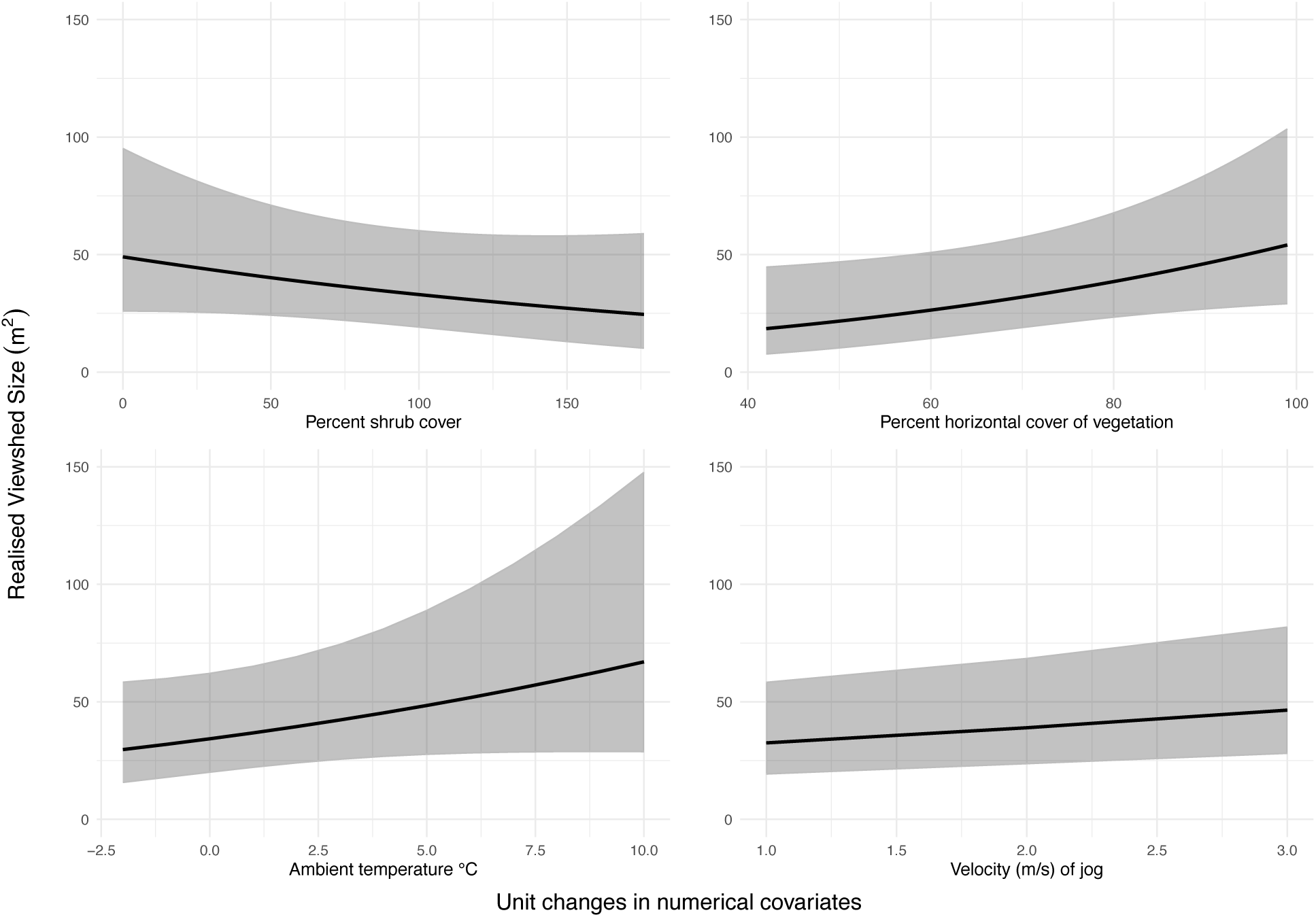
The predicted influence of unit changes of numerical covariates across their measured range in Case Study A, i.e., percent shrub cover, percent horizontal cover of vegetation, ambient air temperate (°C) at time of survey, and velocity (m/s) of each jog, on the Realised Viewshed Size at a given camera trap. For each numerical covariate of interest, all other covariates are held at their mean value for all predictions.

### Case Study B

Our controlled, open field trials suggested that predetermined sensitivity settings have a large, significant influence on the probability of photographic capture and thus RVS (Table 3). Our model estimated a maximum difference of approximately 250 m^2^ in RVS between the lowest and highest sensitivity settings on the Reconyx Hyperfire 2 CTs (Figure 4). The number of photos taken per trigger had a significant negative influence on RVS, where 3 and 5 photographs per capture produced significantly lower RVS than 1 photo (but were not different from each other; Table 3, Figure 4). Jog-tests conducted in the daytime produced significantly higher RVS (^x̅^ = 151 m^2^, SE =122–191 m^2^) than the post-sunset jog-tests (^x̅^ = 73 m^2^, SE = 51–101 m^2^, Table 3, Figure 5).

**Figure 4.**
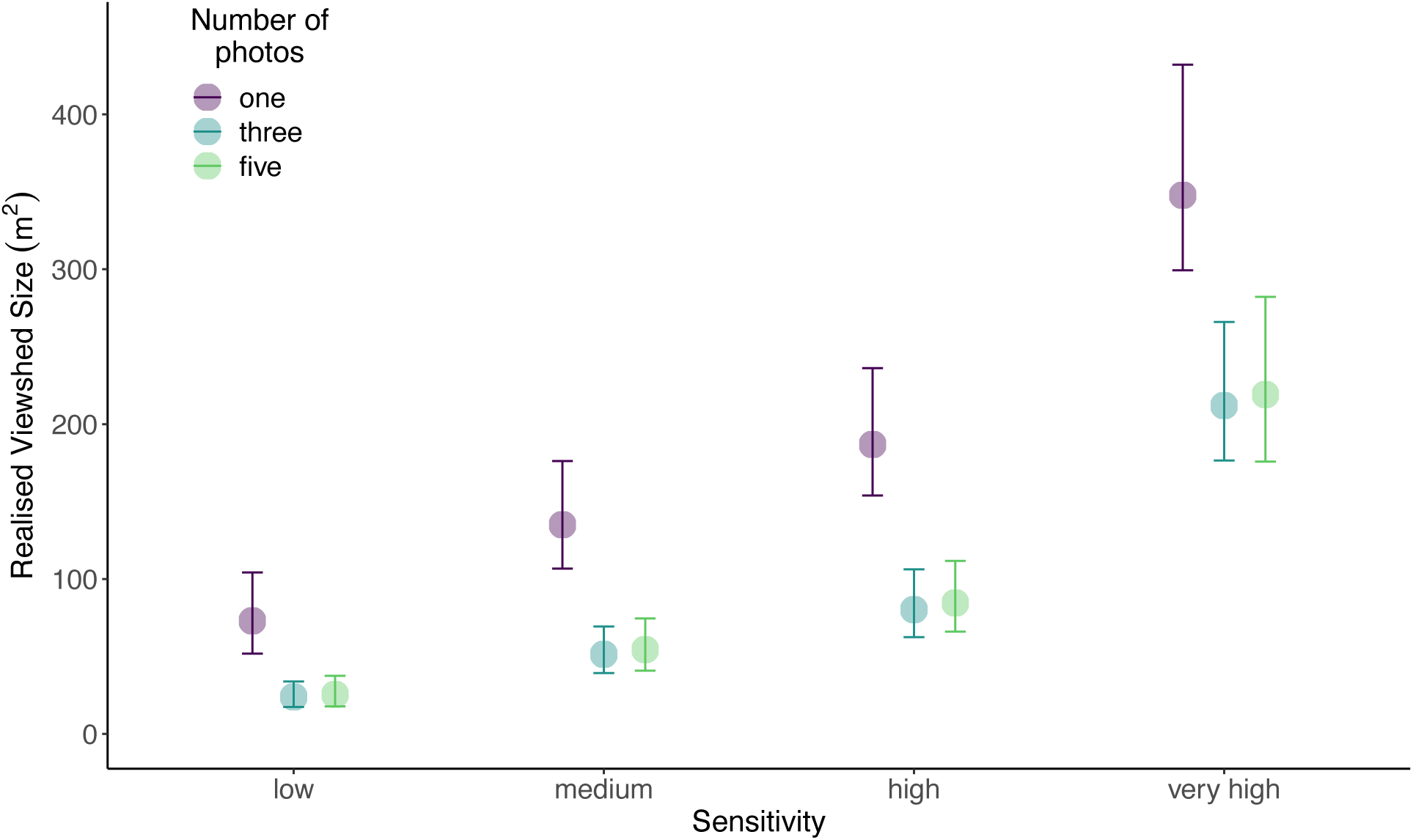
Effects of different sensitivity settings (low, medium, high, very high) and number of photos taken per each Passive Infrared Motion trigger (one, three, five) on the predicted Realised Viewshed Size (m^2^ ± SE) of Reconyx Hyperfire 2 camera traps (*n* = 26) predicted from Generalized Additive Mixed Models during our controlled setting trials for Case Study B.

**Figure 5.**
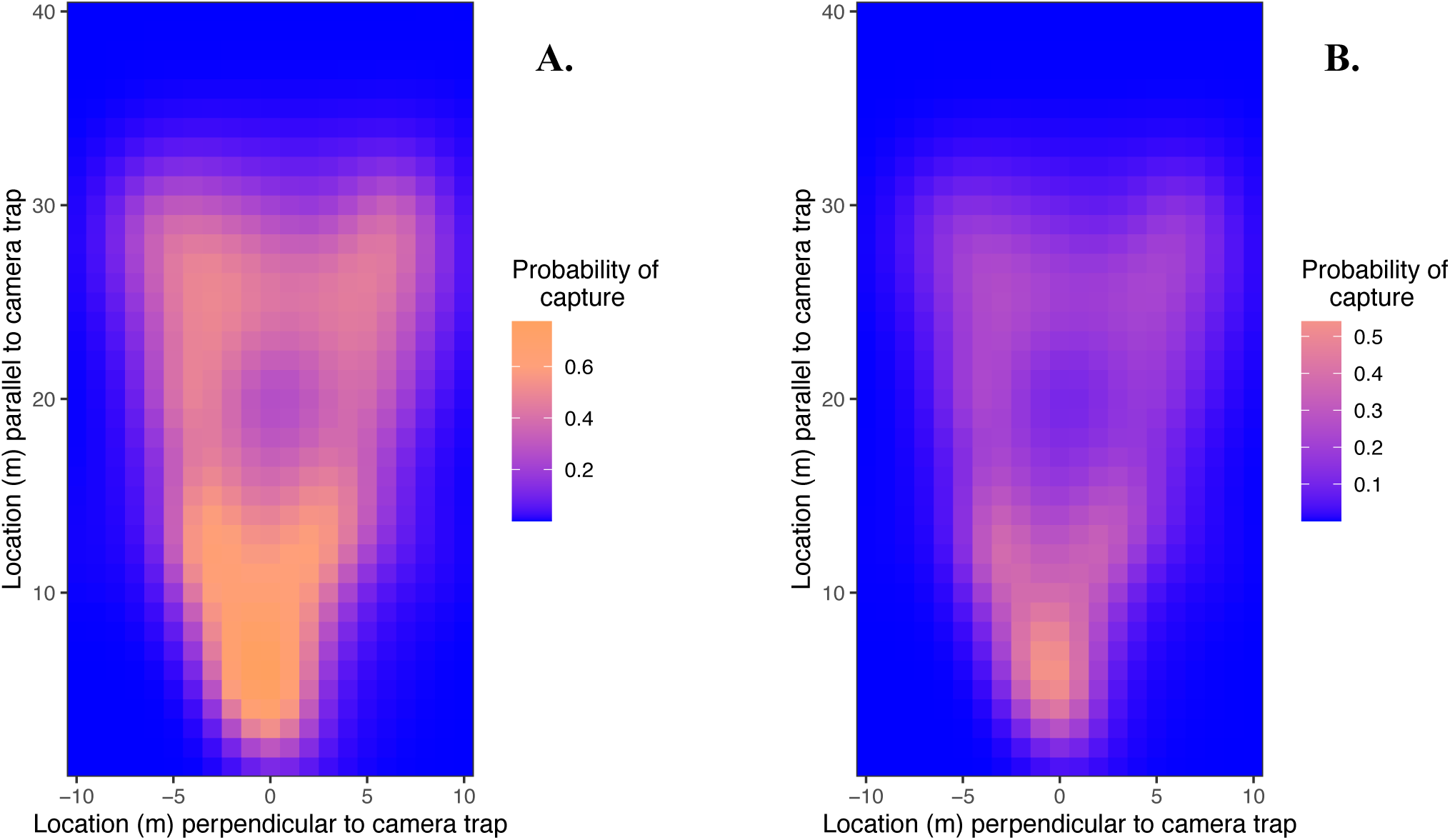
Predicted probabilities of photographic capture, predicted with Generalized Additive Mixed Models, on a 1×1 m grid for Reconyx Hyperfire 2 cameras, assuming a “medium” sensitivity setting, and a single photo taken for each Passive Infrared motion detector trigger during **A**. daytime trials and **B.** nighttime trials. Cameras are located at position [0,0] on each figure. The Realised Viewshed Size (± SE) were determined to be 151 m^2^ (122–191 m^2^) for daytime (**A**) and 73 m^2^ (53–101 m^2^) for nighttime (**B**).

## Discussion

To determine CT viewshed size that is scaled by capture probability and specifically for use in the Random Encounter Staying Time (REST) and Time in Front of Camera (TIFC) density estimation models, we enumerated the Realised Viewshed Size (RVS). The RVS is the theoretical area a CT monitors with 100% capture probability. Here, we showed how the RVS is influenced by local environmental variables and internal CT settings. We determined the RVS through a spatially predictive kernel-based photographic capture probability analysis, providing new insight into how capture probability varies within a CT’s viewshed. Our analysis suggested that capture probabilities and the associated RVS are sensitive metrics and can vary substantially depending on both environmental influences and user-defined settings. Understanding and accurately estimating the RVS is crucial when implementing viewshed-based density estimators and may also contribute how researchers interpret CT-based occupancy.

### Utility for viewshed density estimators

The RVS provides multiple novel considerations for use in some viewshed density estimators. In literature that uses Effective Detection Distance (EDD) with viewshed density estimators (e.g., Becker et al., 2022; Fisher et al., 2023; Palencia et al., 2021), researchers consider periods of time animals were never photographed, but likely present in the viewshed. For example, it is common practice to assume an animal was in a viewshed for all the time in between the first and last photographs of an event. Though this assumption is likely true, it can undoubtedly lead to an overinflation in density estimates (e.g., as seen in Becker et al., 2022; Fisher et al., 2023). Furthermore, it presupposes the animal did not leave then return to the viewshed area. By considering the probability of missed captures our method to estimate RVS removes the need to assume where animals are or are not in the viewshed in the absence of data. Thus, we think the RVS would improve density estimation specifically when used with the REST or TIFC models or derivations (Hogg, 2021; Nakashima et al., 2018; Warbington & Boyce, 2020) that can consider single photographs and their temporal footprint.

### Case Study A

RVS values generated from our Case Study A model are substantially smaller than other published literature. For example, an average Reconyx Ultrafire CT in our study was estimated to monitor an area of 16m^2^ (ranging from 12–22m^2^) with a perfect capture probability. Our average Reconyx RVS, in the Case Study A field trial, was less than 10% of the total possible area the CT advertises (Reconyx, 2022) and still smaller than other published monitoring areas (Becker et al., 2022; Garland et al., 2020). Thus, if implementing the previous detection areas, i.e., 16 m^2^ vs 315 m^2^, on viewshed density estimators, density estimates would differ by an order of magnitude. Although CT capture probability area cannot be unilaterally applied across different studies in varying geography and with different external influences, our work highlights the importance of conducting a standardized test on all CT of a study when attempting to estimate monitoring areas.

In Case Study A, our goal was to develop an accurate metric to estimate the physical space CTs monitor, specifically for use with viewshed density estimators. Though not statistically significant, our environmental covariates, i.e., shrub cover, horizontal cover, temperature, and jog velocity, still contributed considerable variability in RVS. For example, when holding other covariates at mean values, average per-unit changes in vegetation structure, shrub cover and horizontal cover can influence RVS by ∼0.2 m^2^. Although changes are small over single percentage changes, if we consider CTs with 50% differences in either shrub or horizontal cover, RVS could be influenced by over 10 m^2^. Vegetation influences are likely even greater in affecting RVS, considering shrub and horizontal cover are additive, resulting in a compounded effect on RVS.

We detected a relatively small influence of unit changes of temperature on RVS, with a mean of ∼2 m^2^ per °C. The small influence of temperature is intuitive given the short duration when we conducted jog-tests. For example, we conducted our Case Study A jog-tests on the edges of winter (April and November 2023). The time of year allowed us to safely access remote field sites, in addition to providing conditions consistent with when traditional density estimation surveys take place, e.g., aerial flight surveys in late winter, for future applications of our work.

As a result, many of our covariates show little range. Ambient air temperatures at time of survey only ranged from -2–10°C, and total shrub was measured after leaf senescence in the fall, limiting observed variability. Although we would not expect CTs to decrease performance until - 20°C or below (Reconyx, 2022), it is worth noting that other research has observed CTs performing best around 0°C, where higher false-negatives occur in positive temperature ranges (Jacobs & Ausband, 2018), and negative temperature ranges and associated weather can contribute to decreased performance (Maile et al., 2023). Conducting jog-tests during other seasons and within a greater range of forest types and habitat will likely contribute to detecting statistical differences in environmental covariates and help determine if a CT’s RVS varies more substantially across space and time (e.g., as in McIntyre et al., 2020; Moeller et al., 2023; Moll et al., 2020; Sultaire et al., 2023; Urbanek et al., 2019).

Though our jog-tests were conducted at relatively consistent velocities, we detected a large difference in RVS between unit (m/s) changes in jog velocity, with a mean of ∼35 m^2^. Further, our predictions show that slower jog speeds lead to higher RVSs. Our jog velocity results stand in contrast to previous findings (e.g., Del Bosco, 2021) who found that faster jog velocities generally lead to higher capture rates at CTs. The velocity discrepancy may be explained by the habitat in which jog-tests were conducted. Specifically, our tests took place in primary and secondary conifer and mixed forests where as Del Bosco’s (2021) work occurred in more open, sagebrush-mountain habitat. We suggest researchers consider both habitat composition and effective movement speed of the species of interest when implementing the jog-test method.

Cross-validation of the probability of photographic capture occurring in each 1×1-m cell assessed during jog-tests highlighted that our RVS modelling framework had useful predictive accuracy (ROC score of 0.76). In addition, environmental covariates explained substantial variation in our model, as determined through average unit-changes. The variation explained by covariates, along with our cross-validation results, suggest that, in our geography, we can predict RVS well at CTs where we did not conduct jog-tests, but did collect vegetation data.

### Case Study B

Overall, sensitivity settings on Reconyx Hyperfire II CTs greatly influenced the predicted RVS. Differences between lower sensitivity settings were smaller but bme more pronounced as the sensitivity settings became higher. These results show how drastically a CT monitoring zone can change with sensitivity settings, e.g., from ∼75m^2^ to ∼350m^2^ in open conditions, reinforcing the importance of consistency and assessing CTs in each location they are placed. Our results highlight the balance researchers need to consider when altering sensitivity settings to decrease false negative photographic captures, while risking tremendous loss of area in photographable space.

Predicted RVS for trials set to take 3 or 5 photos per trigger were significantly smaller than trials with 1 photo per trigger, but not different from each other. As we take more photographs per event, the CT will be less able to capture future photographs. Failure to capture future events is because, even on rapid fire modes, it takes longer for to capture and write 3 or 5 photographs than a single photograph. When considering refractory period, setting CTs to take more photos will greatly reduce the available time a CT has to take a photo and thus, the RVS. Though additional photos per trigger can be very useful in the identification of species, additional photos have an overall negative influence on a CTs viewshed area. Intuitively, trials conducted in the daytime had significantly higher RVSs compared to post-sunset trials. A lower nighttime RVS is likely because the visible range of the CT will be reduced during nighttime, in addition to a reduced contrast from the subject to the background. Nighttime RVSs will be particularly important on CT studies for species that are predominately nocturnal, as viewshed area may be lower than anticipated.

### Camera refractory period

Our two case studies observed opposite effects of the refractory period variable. In Case Study A, we observed a significant negative relationship between capture probability within a 1×1-m cell and whether a photo was captured in the previous two seconds. The refractory period result met our null expectations that, due to delay, our CTs deployed in the field were less able to capture photos immediately after a previous photo had been taken. Our Case Study B model, however, exhibited the opposite trend and the refractory period variable was significantly positively correlated with capture probability, i.e., photos were more likely to be taken immediately following a previous capture. These contrasting results were likely due to the way we structured the Case Study B trials, where many trials took more than one photo per trigger.

For example, in trials where CTs took 3 or 5 images per trigger, by artifact of CT shutter speed, the time it takes to capture 3 or 5 images is longer than 1 image. Thus, the refractory period of a CT would be longer if the CT is set to capture more images per trigger, e.g., potentially 3–6 seconds after the first photo in the series is captured.

The elongated refractory period caused by trials with more than 1 photograph taken per trigger is also likely responsible for the divergence in capture probability observed in our graphical representation of RVS in Case Study B (Figure 5). During trials where CTs were set to take multiple photos, if captures were registered at the start and end of transects, where a refractory period occurred near the midpoint of the trial, the pattern we observe of decreased capture probability directly perpendicular to CTs past ∼14m could be explained (Figure 5). The gap in photograph captures would not be observed at closer distances to CTs because of CT lens angle—there is physically less space within a CT’s viewshed, explaining the gap in detection probability for trials with multiple photos per activation (Figure 5).

### Limitations and future directions

The functioning and performance of CTs remains an understudied facet of the field that certainly influences CT analyses and thus interpretation of CT-based research. In our work, how PIR sensors perform across temperature and seasonal gradients is particularly important. PIR detectors in CTs function by sensing a difference in temperature of a subject from ambient temperature, in addition to movement, to capture a photograph (Reconyx, 2022; Welbourne et al., 2016). The interacting effects of ambient temperature and seasonality will influence the amount of heat wildlife subjects will emit. For example, in winter, ungulates have thick winter coats that allow them to retain heat and survive harsh conditions and as such, less heat may be escaping the coat (Parker and Robbins 2018). A lack of heat emission has been previously observed in thermal imaging studies, where thermal CTs have a difficult time picking up certain species during certain temperature ranges (Kuhn & Meyer, 2009; Zabel et al., 2023). In addition, the effects of snow cover in northern geography and how it may contribute to capture of animals, through heat recognition or increased exposure merits further research. As a result, capture probability may be even lower during mid-winter months for some wildlife species.

CTs are influenced, to some degree, by the size of the subject they are capturing (DeWitt & Cocksedge, 2023). Other viewshed size estimators (e.g., Effective Detection Distance; EDD) have found substantial variation in the EDDs between different sized animals (Hofmeester et al., 2017; Becker et al., 2022). Because EDD measures the minimum distance where at least one photograph of a species is captured, we think size may be more influential than the RVS—where we measure the probability a photograph will be captured given a subject is in a discrete space. For example, Urbanek et al. (2019) observed that raw number of photographs at CTs were similar for species groups that were generally the same size. The pattern in Urbanek et al. (2019) data suggests a potential trigger threshold where above certain body sizes, the number of photographs taken (and thus RVS) may increase. Thus, we think that our approach will work well for some species, particularly ungulates, based on height, however, we would not recommend our methods when applied to species with large size discrepancies, particularly smaller, to humans.

Our jog-test trials used linear transects at an angle perpendicular to CTs optical axis. Such linear transects may influence RVS values. In realistic settings, wildlife may move simultaneously along both perpendicular and parallel angles relative to a CT’s optical axis. In addition, a subject’s body orientation, and thus size, may change as the subject moves throughout a CT’s viewshed. Effects on capture probability and RVS would be non-linear and dependent on each specific capture event being photographed. Despite our linear transects, we investigated the probability of photographic capture given a subject was in a delineated location and fit those probabilities as a kriged surface using a Gaussian process spline. Thus, because of our statistical approach we think the influence of other angles of approach within a CT viewshed on the RVS would be negligible.

Our GAMM model was developed to incorporate important predictor variables specifically at our field site in Riding Mountain National Park (MB, Canada). We hypothesized the largest influencing factor to CTs would be vegetation, specifically shrub cover. Although vegetation cover in our study had a relatively lower influence on RVS (Table 3, Figure 3), the importance of different predictor variables, will likely depend on the geography and habitat conditions of the study. For example, topography (Sultaire et al., 2023) and weather (Madsen et al., 2020) have been found to influence CT capture probabilities in some geographies. Thus, if implementing our method, researchers should consider all potential important predictor variables that may influence capture probability at their study sites.

## Conclusion

We use a standardized, *a priori* field and analytical protocol to predict the variable probability of photographic capture at CTs while incorporating internal and environmental influences on CT performance. We use our predictions of capture probability to estimate the Realised Viewshed S1ize (RVS), a novel estimator that is the scaled area in front of CTs that represents a 100% capture probability. Our results highlight how variable CT performance can be and provide a framework for researchers and other CT users to account for variable viewshed areas and capture probabilities for some taxa of wildlife. RVS may contribute to increasing the reliability and precision of CT-based density estimators and thus forward the growing branch of CT research.

## Data availability

Data and code used in this analysis will be made available at the link following publication: https://github.com/wildlifeevoeco/RealisedViewshedSize

## Notes

### Competing Interest Statement

The authors have declared no competing interest.

### Summary of Updates

Figure 3 has been revised following an error noticed in the prediction function used. We now show what we intended to, and what is described in text.

